# Using a Bayesian network model to predict effects of pesticides on aquatic community endpoints in a rice field – A southern European case study

**DOI:** 10.1101/2022.10.19.512688

**Authors:** Sophie Mentzel, Claudia Martínez-Megías, Merete Grung, Andreu Rico, Knut Erik Tollefsen, Paul J. Van den Brink, S. Jannicke Moe

## Abstract

In recent years, Bayesian network (BN) models have become more popular as a tool to support probabilistic environmental risk assessments (ERA). They can better account for and communicate uncertainty compared to the deterministic approaches currently used in traditional ERA. In this study, we used the BN as a meta-model to predict the potential effect of various pesticides on different biological levels in the aquatic ecosystem. The meta-model links the inputs and outputs of a process-based exposure model (RICEWQ), that is run with various scenarios combination built on meteorological, hydrological, and agricultural scenarios, and a probabilistic case-based effect model (PERPEST), which bases its prediction on a database of microcosm and mesocosm experiments. The research focused on the pesticide exposure in rice fields surrounding a Spanish Natural Park, considering three selected pesticides for this case study: acetamiprid (insecticide), MCPA (herbicide), and azoxystrobin (fungicide). For each of the pesticide types, the developed BN model enabled the prediction of their effects on biological endpoints, endpoint groups, and community in an aquatic ecosystem. Also, it enables comparison between the different pesticide types, their effects on endpoint groups and community. While directly linking future scenarios of climate and agricultural practice to the exposure concentration and indirectly linking them to the effect on biological endpoints as well as community. In summary, azoxystrobin and MCPA seem to have a higher predicted risk for the community with at least one of the biological endpoint being effected compared to acetamiprid. Generally, the developed approach facilitates the communication of uncertainties associated with the predicted effect on different biological levels of the aquatic ecosystem. This transparency in all model components can aid risk management and decision making.

## 1 Introduction

In the future, changes in agricultural practices, as for instance, the use of new or more plant protection products (Delcour et al., 2015) can cause a change of risk to biodiversity in aquatic ecosystem. Some agricultural methods lead to an intensive use of pesticides. This is of special concern in the Albufera Natural Park (Valencia, Spain) a lake enclosed by rice fields, known for its diversity in bird and fish species (Soria, 2006). In this area, the rice production and other anthropogenic stressors already had a negative impact on lake’s water quality and ecosystem health throughout the last century (Calvo et al., 2021; Vera-Herrera et al., 2021). Realistic assessment of risk posed by these expected stressors is therefore crucial for the future ecological sustainability of the Albufera National Park.

### 1.1 Importance of effect assessment using predictive models

Today’s ERA is mostly based on deterministic approaches, usually relying on single value risk estimation, such as risk quotients, to provide predictions based on individual effects data (for larger organisms) or sub-populations (for smaller organisms run in lab tests with subsets of a population). Thus far, good and reliable assessment of pesticide risk requires realistic exposure and effect data, as well as understanding of ecosystem processes (Schmolke et al., 2010).

ERA often uses monitoring studies for exposure assessment, though these can be time-consuming and expensive, and their results are quite site specific and have a wide range of uncertainties (Lammoglia et al., 2018). Ergo, for pesticide ERA using pesticide fate simulation models (Pereira et al., 2017) is a needed tool for the prediction of exposure concentration and for the characterization of spatial and temporal long-term patterns. Moreover, as future land-use and climate changes (CC) are expected to alter the distribution and fate of pesticides in the aquatic environments. In the Mediteranean, it is expected that droughts occur more frequently, and water is less available, thereby resulting in lower dilution. On the other hand, severe precipitation events are expected to occur more often which may result in higher pesticide runoff. For this southern region, an expected increase in temperatures, may facilitate microbial degradation of pesticides but also higher uptake by organisms (Arenas-Sánchez et al., 2016; Balbus et al., 2013; García de Jalón et al., 2014; Noyes et al., 2009). In recent years, the CC’s influence on pesticide fate and transport has been the subject of increased concern (Bloomfield et al., 2006; Delpla et al., 2009; Lamon et al., 2009; Noyes et al., 2009). Exposure prediction models can aid exposure assessment in cases where monitoring data is scarce, or for example assist the analysis of future land-use and CC impact in prospective exposure assessment. As these process-based exposure models are able to integrate a wide diversity of scenario combinations such as agricultural practices, soil properties, crop types and meteorological conditions, consequently being a relative rapid and cost efficient tool assessing the exposure of pesticides to the environment (Lammoglia et al., 2018).

For effect assessment based on toxicity test, indirect effects are frequently not considered, nor is the complexity of the population and population dynamics accounted for neither is the complex interactions occurring between populations in a community structure. Whilst some exceptions may be mesocosm studies, that are often based on single-chemical and single-species, certain environmental media (soil, water, or sediment) and under laboratory conditions (Di Guardo & Hermens, 2013). The traditional assessment, species response and interaction have to be extrapolated and accounted for by applying assessment factors to the most sensitive toxicity test or hazard concentration from a species sensitivity distribution (Schmolke et al., 2010; Topping et al., 2020). The current effect assessment is lacking insights on the concentration-response relationship between different trophic levels of the ecosystem (Van den Brink et al., 2006). Furthermore, ERA needs to better consider the interaction of contaminant and noncontaminant stressor (Landis et al., 2013), such as changes in climate conditions (temperature & precipitation) as well as changes in land use practices as they can lead to shifts in ecosystems, their hydrological processes. In turn, this may lead to changing responses to contaminants by affected species (Landis et al., 2013). Multiple stressors have been found to affect freshwater ecosystem functioning and structure long term, and can influence resilience and recovery of the ecosystem (Polazzo et al., 2022). Instead of basing the effect assessment only on single species toxicity tests, multi species models can be used to predict and analyse possible indirect effects within community. Food chain models can include food-web models, that only consider trophic relationships, or community models, that also consider some inter-species interaction (Larras et al., 2022).

Some multi species models are based on case-based reasoning (CBR). These CBR based models are based on *“a paradigm of artificial intelligence and cognitive science that models the reasoning process as primarily memory based. Case-based reasoners solve new problems by retrieving stored ‘cases’ describing similar prior problem-solving episodes and adapting their solutions to fit new needs”* (Leake, 2001). They can consider various factors such as endpoints, experimental ecosystems and test design in their prediction. One such model is the PERPEST model used in this study (Van den Brink et al., 2002), it predicts direct effects on communities while accounting for some indirect effect and interaction among species groups informed by observations from mesocosm studies (Davis et al., 2013) while considering the mode of action in its prediction (Larras et al., 2022; Van den Brink et al., 2006; Van den Brink et al., 2002).

### 1.2 Probabilistic environmental risk assessment method needed to handle sources of variability and uncertainty

Often, European prospective ERA is based on toxicity exposure ratios or other single values where potential risk is often calculate by comparing predicted exposure (concentration) to no-effect concentration (Di Guardo & Hermens, 2013; Schmolke et al., 2010). This deterministic approaches of ERA describe risk as a “margin of safety” using uncertainty factors, or the exceedance and frequency of exceedance of safe thresholds, to derive qualitative output that lacks indication of the level of certainty related to the input and output parameters (EUFRAM, 2006). In reality however, pesticide exposure and effects have spatial and temporal variability, and are determined by environmental and biological characteristics, as well as pesticide application pattern (FOCUS, 2007). Improving prospective ERA, and by considering and integrating future scenarios in the prediction risk of to the aquatic environment would improve prevention further and future damage (Topping et al., 2020). Some of the limitations of traditional ERA can be overcome with probabilistic approaches that characterize both toxicity and exposure, typically using distributions or assigning probabilities. Consequently, they are able to account for variability and uncertainty better (Carriger & Newman, 2012; EUFRAM, 2006; Solomon et al., 2000; Verdonck, 2003). The use of probabilistic approaches has also been recommended by the European Union (Jager et al., 2001). Commonly used probabilistic methods are joint probability curves, quantitative overlap, or risk quotient distribution (Campbell et al., 2000; Mentzel et al., 2021; Verdonck, 2003). These commonly used probabilistic approaches outputs can be hard to understand and communicate to decision-makers (Dreier et al., 2020; Giddings et al., 2000).

Bayesian networks can overcome some of these limitation and better communicate and quantify uncertainties to decision-makers and other stakeholders (Carriger et al., 2016; Carriger & Newman, 2012). While being used in situations where data is limited or processes lack characterization, BNs are able to incorporate these various sources of information e.g., expert elicitation, model outputs or literature (Carriger et al., 2016; Carriger & Newman, 2012; Gibert et al., 2018; Hamilton & Pollino, 2012). Besides, BNs have the ability to act as a meta-model (e.g. Mentzel et al. (2022)), allowing for the incorporation of inputs and outputs from various different models (in a single model). Summarized, they are probabilistic graphical models that contain nodes (variables) linked through arcs representing conditional probability tables (CPT) (Aguilera et al., 2011; Kaikkonen et al., 2021). The nodes have assigned states (intervals) that can be quantified by probabilities and probability distributions. Based on new evidence, these Direct Acyclic Graphs (DAG) uses Bayes’ rule to update the probability distributions throughout the network (Carriger et al., 2016; Kanes et al., 2017). Overall objective, is to predict risk of pesticides to biological communities represented by multiple biological species groups. To achieve this, the probability of effect on biological endpoints was predicted for the different peticide. Secondly, the developed BNs predicted the cumulative probability of effects of a pesticide on different endpoint groups (invertebrates etc.) as well as the whole community. Thirdly, the effects of different pesticide types on different endpoint groups are compared as well as community level. Lastly, this BN model aims to predict the probability of effects under scenarios of pesticide application or climate change.

## 2 Material and methods

### 2.1 Description of the case study region

The study area is a coastal wetland around five kilometres south of Valencia on the Mediterranean Spanish coast, with an area of about 210 km^2^ (Figure 1). The Natural Park has ecological relevance as it is a nesting and transfer point for approximately 250 species of migrating birds and mentioned in as a special protection area by Birds Directive (Directive 2009/147/EC), listed as European habitat in Natura 2000, and Ramsar Convention of wetlands (Calvo et al., 2021; GV, 2020; Vera-Herrera et al., 2021). Within its bound, 34% of Spanish rice is produced (Canet et al., 2003) as 73% of the wetland is dedicated to rice cultivation (Vera-Herrera et al., 2021). Also, the lakes water level is regulated by a network of irrigation channels and seasonal rainfall (mainly spring and autumn). Agricultural and other anthropogenic activities had negative impact on the shallow (1-1.2 m mean depth) and oligohaline (1-2% salinity) lake located in the centre of the wetland (Calvo et al., 2021; Vera-Herrera et al., 2021).

**Figure 1.**
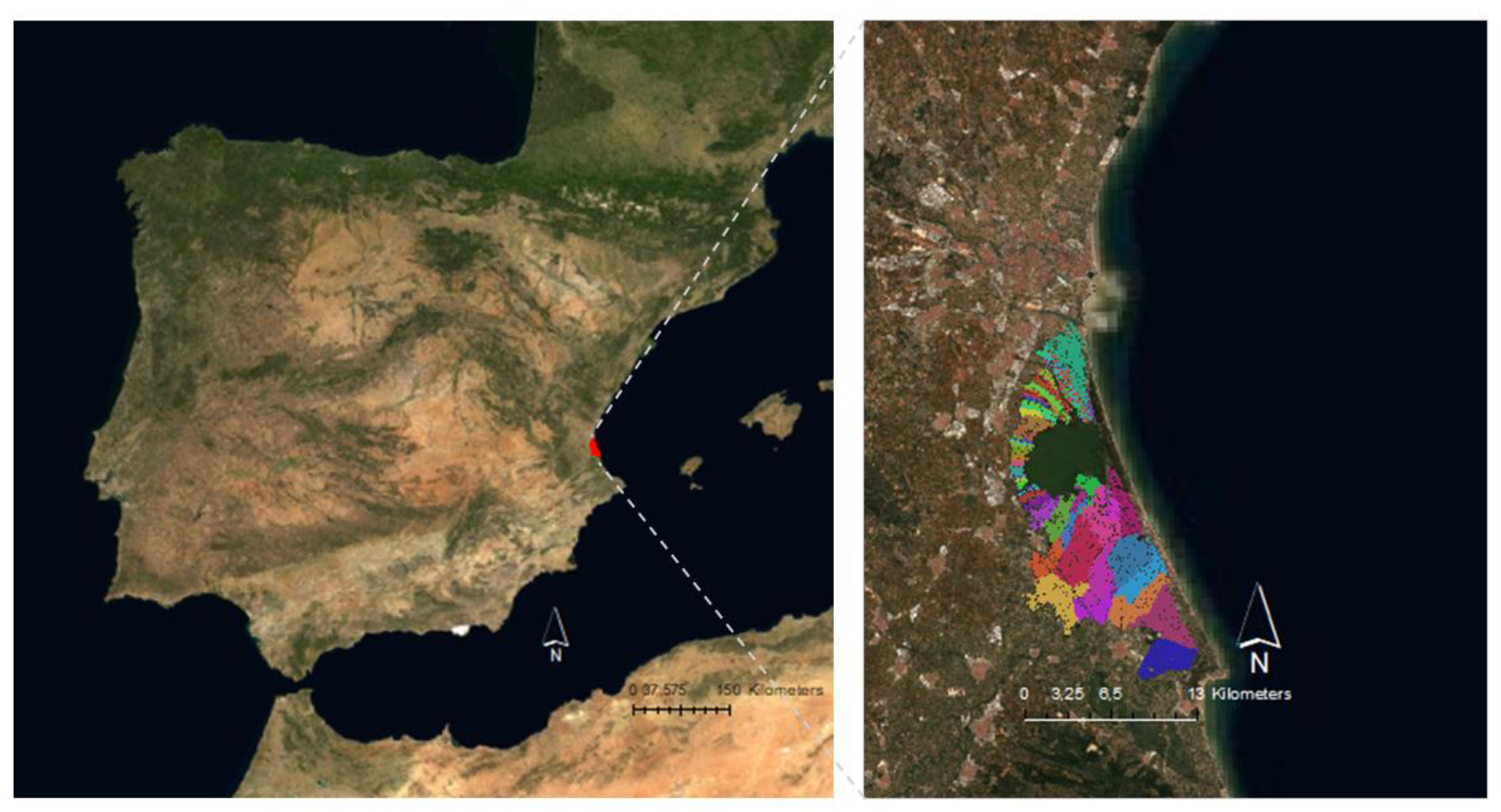
Location of Albufera Natural park near Valencia (red) and location of rice field clusters (coloured areas) (adapted from IGN (2022), Retrieved from www.ign.es, Accessed on 28 April 2022)

### 2.2 Bayesian network conceptual model and assumptions

The general approach is to integrate predicted outputs from both exposure and effect prediction models into a Bayesian network serving as a meta-model. In this study, the RICE Water Quality model – RICEWQ was used to simulate the pesticide exposure in the water in rice paddies (Karpouzas & Capri, 2006; Miao et al., 2004) and the Predicts the Ecological Risks of PESTicides (PERPEST) model was used to simulate the pesticide effect to various biological endpoint (Van den Brink et al., 2002). We developed a BN meta-model structure incorporating temporal variability in the effect estimation of pesticide for various endpoints in the aquatic ecosystem. Thus, the simulated peak concentrations (RICEWQ output) are converted to a probability distributions and the gradients predicted for each biological endpoint (PERPEST outputs) are manually added as prior probabilities in the conditional probabilty tables of the related biological endpoint nodes in the BN.

The BN model is composed of three modules: the scenarios and exposure module (blue), effect on biological endpoint module (green) and effect on community (grey) module (Figure S. 3). The first module, scenario and exposure, is composed of the scenario combination (red) that define the exposure concentration distributions fitted to the RICEWQ model output. The second module is derived by the PERPEST model output that provides the effect concentration states and the prior probabilities of the biological endpoint nodes. In the third module “cumulative risk to community”, each of the biological endpoint nodes are transferred to Boolean nodes (true/false) before being aggregated to their respective endpoint group nodes (light grey) (e.g. effect on Vertebrates) and further aggregated to the community level (Figure 2). Thus the node “Macrophytes bool” is meant to quantify the probability of a pesticide effect to macrophytes (true/false); the node “Effect on plants” will quantify the cumulative probability of effects to any of the plant endpoints; and the node “Effect on Community” will quantify the cumulative probability of effects to any of the endpoint groups.

**Figure 2.**
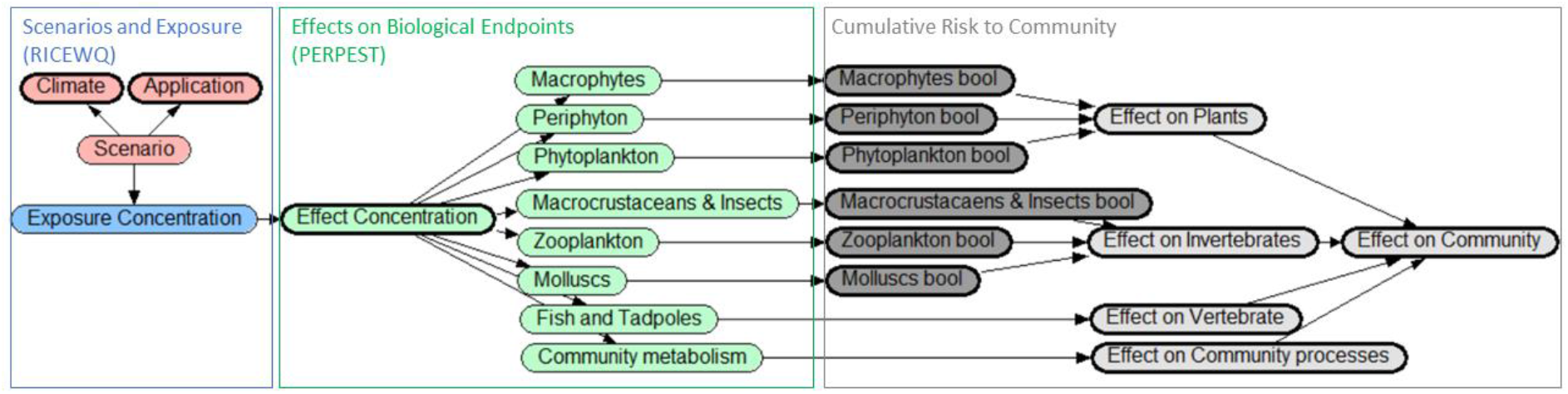
Conceptual model for the effect estimation of a pesticide (acetamiprid) on an aquatic community. Pesticide exposure derives input from the RICEWQ model and is determined by the associated future and application scenarios. The PERPEST model input derives the effect on biological endpoints and endpoint groups and in turn the effect on the community.

In this study, the BN was constructed with the Netica software (Norsys Software Corp., www.norsys.com). For each of the selected pesticides, the BN illustrates the predicted exposure concentration and effect on the various biological endpoint and endpoint groups and summarizing effect on community. The parameterized model can be run by selecting a set of scenarios e.g. climate time and application scenario as evidence. The probability distributions will then be updated throughout the BN to predict the probability of effect classes on the output nodes. Model assumptions and a more detailed node description are detailed in Table 1.

**Table 1.** Bayesian network node description containing the node name, type, number of states and information source.

| Module | Node name | Node type | No. of states | Node input source |
| --- | --- | --- | --- | --- |
| Scenario and Exposure | Climate time | Categories | 3 | Scenarios (2008, 2050, 2100) |
|  | Application scenario | Categories | 2 | Scenarios (baseline, baseline+50%) |
|  | Scenario combination | Categories | 6 | Combination of the scenarios |
|  | Exposure concentration | Intervals | 8 | Scenario-dependent distribution: Normal Distribution (mean, sd) |
| Effects on biological endpoints | Effect concentration | Intervals | 6-10 | = exposure concentration with discretization adapted to intervals used by PERPEST |
|  | Biological endpoint node | Ranked categories | 3 | Pesticide effect (no, slight, clear) on biological endpoints as predicted by PERPEST |
| Cumulative risk to community | Effect on endpoints | Boolean | 2 | FALSE = no effect<br>TRUE = slight effect + clear effect |
| | Effect on endpoint group | Boolean | 2 | Effect on functional groups = $1-(1-\text{node}_a) * (1-\text{node}_b) * \dots (1-\text{node}_n)$ |
| | Effect on Community | Boolean | 2 | Effect on Community = $1-(1-\text{node}_a) * (1-\text{node}_b) * \dots (1-\text{node}_n)$ |

### 2.3 Exposure prediction with RICEWQ - prediction and settings

As previously mentioned, the RICEWQ model was developed to simulate pesticide exposure in water of rice paddies. It is a process-based model that at field level simulates pesticide runoff specific for use in rice paddies (Williams et al., 1999). Thus far, it is considered to be most suitable and reliable for higher-tier pesticide fate and exposure prediction (Daam et al., 2013; Karpouzas & Capri, 2006; MED-Rice, 2003). Besides, it has been widely applied in the US (Karpouzas & Capri, 2006; Miao et al., 2004) to track the fate of both parent and metabolite chemicals (Christen et al., 2006). Various processes are reflected in RICEWQ modelling such as biological, hydrological and physicochemical processes (Wang et al., 2019).

This exposure prediction model requires the following inputs: daily weather information, paddy soil properties, pesticide chemical properties, pesticide management information, and water management practices (Wang et al., 2019). The reader is referred to (Williams et al., 1999) for a more detailed information of the model function, assumptions and description.

In this study, the latest version, RICEWQ 1.92 (Waterborne Environmental Inc, 2022) was run for various scenarios incoprorating different rice crop types, management practices, meterological and hydrological conditions, and for selected pesticides used for the rice crops in the region. Moreover, we selected three active substances that are regularly applied by farmers around Albufera lake. The derived pesticide application scenario is based on the recommended manufacturer dosages from which we derived two scenarios: one maximum recommended dosage (referred to as Baseline application throughout this study) and one that is 150% of that baseline dosage (Baseline+50%). Initially we had aimed to have at least 3 different emission scenarios’ climate projections to include more variability and used them as input for the RICEWQ model runs, based on what had been previously used in a study by Pool et al. (2021). Though, available prediction data was limited for the specific meteorological station 8416 located near the National park. Therefore, only one climate prediction data set was collected from AEMET (2021) at “Climate projections for the XXI Century – Daily data”, derived with the model “GCM MPI-ESM-LR” and an emission scenario “representative concentration pathways (RCP) 8.5”. Based on this data set, three “climate-time” scenarios for the years 2008, 2050 and 2100 were used to run the exposure prediction model.

The exposure prediction model was run for 552 rice crop clusters. The maximum exposure concentration from each cluster was used to fit to the exposure distribution. A detailed description of the assumptions made to derive these clusters, as well as the automatization of the RICEWQ with a handy interface are available in Martínez-Megías et al. ([in prep]). In total six different scenarios were developed each of which is the combination of year and application scenario, and pesticide, and run with autoRICEWQ, (open source under GPL-3.0 License, programmed in Python 3) and can be accessed at Fuentes-Edfuf and Martínez-Megías (2022).

The prior probability of exposure concentration node is assumed to be a normal distribution with varying mean and standard deviation depending on the scenario combination (Table 2). Onwards in this paper the focus will be on the predicted effect of 2050 as it was considered sufficient for the concept development. The predictions from the other years will be presented in supplementary, as for valid predictions of the effect of climate models on the various biological endpoints, more climate model scenarios would be needed to account for uncertainty and variability in future.

**Table 2.** The exposure peak concentration means and standard deviations used as input on the Bayesian network for the selected pesticides and scenarios, also detailing the year and application scenario. (three significant digits were chosen)

| Scenario | Pesticide type | Pesticide | Year | Application | Mean (µg/L) | Sd (µg/L) |
| --- | --- | --- | --- | --- | --- | --- |
| 1 | Insecticide | Acetamiprid | 2008 | Baseline | 0.35 | 0.27 |
| 2 |  |  |  | Baseline+50% | 0.47 | 0.36 |
| 3 |  |  | 2050 | Baseline | 0.86 | 0.45 |
| 4 |  |  |  | Baseline+50% | **1.14** | 0.60 |
| 5 |  |  | 2100 | Baseline | 0.58 | 0.20 |
| 6 |  |  |  | Baseline+50% | 0.77 | 0.26 |
| 1 | Fungicide | Azoxystrobin | 2008 | Baseline | 80.5 | 10.0 |
| 2 |  |  |  | Baseline+50% | **121** | 15.0 |
| 3 |  |  | 2050 | Baseline | 71.9 | 9.80 |
| 4 |  |  |  | Baseline+50% | 108 | 14.7 |
| 5 |  |  | 2100 | Baseline | 69.6 | 8.74 |
| 6 |  |  |  | Baseline+50% | 105 | 13.1 |
| 1 | Herbicide | MCPA | 2008 | Baseline | 37.7 | 5.59 |
| 2 |  |  |  | Baseline+50% | **56.5** | 8.40 |
| 3 |  |  | 2050 | Baseline | 33.1 | 5.20 |
| 4 |  |  |  | Baseline+50% | 49.6 | 7.81 |
| 5 |  |  | 2100 | Baseline | 24.3 | 4.79 |
| 6 |  |  |  | Baseline+50% | 36.5 | 7.16 |

### 2.4 Effect prediction with PERPEST – model assumptions and prediction

The PERPEST model was developed to simulate pesticide effects to various biological endpoint (Van den Brink et al., 2002) and can be used for risk assessment of single and mixed application of pesticide (Rämö et al., 2018). It is considered more comprehensive compared to the traditional ERA that uses risk or hazard quotient approaches (Polidoro & Morra, 2016; Rämö et al., 2018). This effect prediction model applies case-based reasoning approach (CBR) to draw empirical ecotoxicological data from micro- and mesocosm experiments (Davis et al., 2013; Rämö et al., 2018). It can predict direct and indirect effects of contaminants while incorporating hydrological properties and acute and chronic exposure in the prediction (Van den Brink et al., 2002). The PERPEST model compares environmental exposure concentrations to previous observations in mesocosm and microcosm toxicity tests to estimate the probability of the pesticide having a toxic effect on various pesticide type dependent biological endpoints and endpoint groups.

The PERPEST model predicts a probability gradient for three (default) effect classes on biological endpoints depending on the modelled pesticide type. Following Van den Brink et al. (2002) these three classes are:

- “No effect” - No consistent adverse effects are observed as a result of the treatment. Observed differences between treated test systems and controls do not show a clear causality;
- “Slight effect” - Confined responses of sensitive endpoints (e.g. partial reduction in abundance). Effects observed on individual sampling dates only and/or of a very short duration directly after treatment; and
- “Clear effect” – severe reductions of sensitive taxa over a sequence of sampling dates, and are demonstrated, but duration of the study is too short to demonstrate complete recovery within 8 weeks after the last treatment (Davis et al., 2013).

The reader is referred to Van den Brink et al. (2002) for more detailed information of the model function, assumptions and description.

In this study, we used the PERPEST model to predict the effect of a fungicide, herbicide and insecticide on the biological endpoint associated with being affected by the different types of pesticides. The selected pesticides for this study were not currently available in the PERPEST case base, therefore their physico-chemical properties were collected from literature and databases such as PPDB (Lewis et al., 2016), PubChem (Kim et al., 2020) and CompTox (Williams et al., 2017). The median hazard concentration (HC50) was calculated for each of the pesticides using MOSAIC (Charles et al., 2017) with EC50 toxicity data collected from ECOTOXicology Knowledgebase (Olker et al., 2022). The used input information that are compared to the toxicity dataset by the PERPEST model are shown in Table 3.

**Table 3.** Chemical, biological and physical properties of the selected pesticides included in the PERPEST model.

| CAS | Chemical name | Type of substance | Mode of action | Molecule group | DT50 (days) | Henry's law (Pa m<sup>3</sup> mol<sup>-1</sup>) | HC50 (µg/L) | Koc (L/kg) |
| --- | --- | --- | --- | --- | --- | --- | --- | --- |
| 135410-20-7 | Acetamiprid | Insecticide | Other insecticide | Neonicotinoid | 2950* | 5.3E-08 | 93 | 199.5 |
| 131860-33-8 | Azoxystrobin | Fungicide | Other fungicide | Strobilurin-quinone outside inhibitor | 6.1 | 7.4E-09 | 503 | 3320 |
| 94-74-6 | MCPA | Herbicide | Other herbicide | Aryloxyalkanoic acid | 19.5 | 0.000055 | 5300 | 56 |

\* above max range - the value was higher than maximum amount of days that could be entered, in this case the max of 1000 days was used.

The PERPEST model predicts the probability no effect, slight effect and clear effect along a gradient of pesticide exposure concentrations for a set of biological endpoints (e.g. insects or phytoplankton). The selection of endpoints depended on the pesticide types (for a more complete description of endpoints, see Supplement Information I Figure S. 1).

The latest version of the PERPEST software (Van den Brink et al. [2002]; version 4.0.0) was used to predict the probability of effects for the three selected pesticides (www.perpest.wur.nl) in this study. In principle, the model was used with default settings, additionally weighted with “toxic unit (TU)” and “DT50”, and exposure being set to “not used”.

An example of the PERPEST gradient output is added in the supplement information (Supplement Information I Figure S. 2), a detailed overview of the output tables used as conditional probability tables of the biological endpoint nodes can be found in the supplement information II. In this example, the effect gradient is displayed for predicted effect of the insecticide (acetamiprid) on the of algae and macrophytes group.

## 3 Results

A parameterized example of the BN is shown in Figure 3 for the insecticide acetamiprid. It displays the event of scenario 4, so for current climate in 2050 and a Baseline+50% application, resulting in the displayed exposure distribution (exposure concentration node). For this event the predicted effect on the biological endpoint varies, with no effect for fish and other macroinvertebrates, and mostly no effect with a probability of 80-90% for rotifers, community metabolism, as well as algae and macrophytes. The highest probability of there to be a clear effect is predicted for macrocrustacea with about 24%. The effect on the endpoint groups also vary, there being no predicted biological endpoint effected in the vertebrates, and most likely non effected in the endpoint group of plants (ca. 97%). The summarizing node effect on community predicts there being an effect on ate least one of the biological endpoints with about 77%.

**Figure 3.**
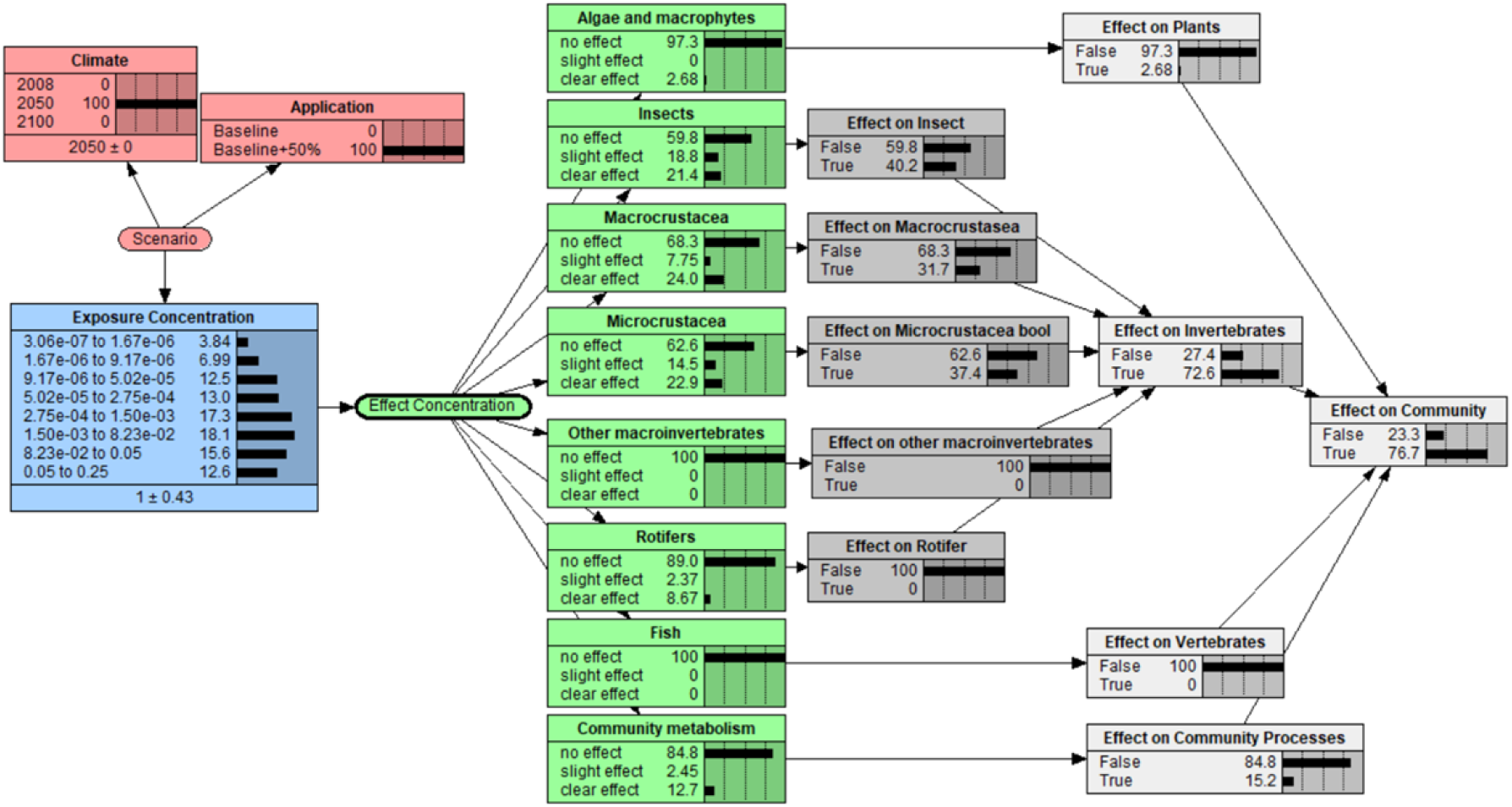
Example of the parameterized Bayesian network for the insecticide acetamiprid. It displays the predicted effect on the biological endpoints and endpoint groups for climate conditions of 2050 and a baseline+50% application scenario.

In the subsequent, the probabilities of the output nodes predicted by the BN are displayed in a bar chart to enable easier comparison between the different scenarios, biological endpoint and pesticides types.

### 3.1 Predicted effect on biological endpoints

Focusing on the biological endpoints, the BN predicted the effect on the insecticides acetamiprid for eight biological endpoints. Insects, macro- and microcrustaceans had a probability up to 30% to be in the state of slight to clear effect. Community metabolism, algae and rotifers were mostly unaffected. fish and macroinvertebrates were most likely to not be affected by the insecticide (Supplement Information I Figure S. 5).

For this fungicide azoxystrobin, eleven biological endpoint were considered by the PERPEST model. Macroinvertebrates, microcrustacean and other zooplankton taxa were the biological endpoints predicted to be clearly affected with a likelihood of 50%, followed by Other zooplankton, phytoplankton, community metabolism, and macrocrustacea. Fish and macrophytes were similarly affected with a probability to be of 15 to 20% to be the “no effect” state. Decomposition and periphytic were predicted to not be affected (approx. 100%) (Supplement Information I Figure S. 7).

The PERPEST model considered eight biological endpoints for this herbicide MCPA. Here zooplankton even had a probability of more than 50% be in the clear effect state and phytoplankton and periphytic had a probability of 25%. Macrophytes and community metabolism were also mostly unaffected, and fish and molluscs were predicted to be not be affected with about 100% (Supplement Information I Figure S. 9).

There were few biological endpoints all pesticides had in common, one of them was macrocrustacea (see Figure S.5, Figure S.7, Figure S.9). For all pesticides the distribution of probabilities over the three states were similar. The fungicide and insecticide were predicted to have a probability of 25% to be in the clear effect state in 2050. Compared to the other pesticides, the insecticide had lowest predicted probability for it to be in a clear effect state with about 20%. All of the pesticides showed the highest probability of the macrocrustacea not to be affected.

### 3.2 Aggregation of the predicted effect from biological endpoints to endpoint groups

The BN model output could be defined for the effect on a specific biological endpoint (Figure 4). In the following an example of acetamiprid is shown that dispalys the aggregation from the PERPEST defined states to the effect on the biological endpoint. It was expected that insects were affected with a probability of approx. 22 %, slightly affected with 18 %, and not be affected with 60 % by acetamiprid (Figure 4a).

**Figure 4.**
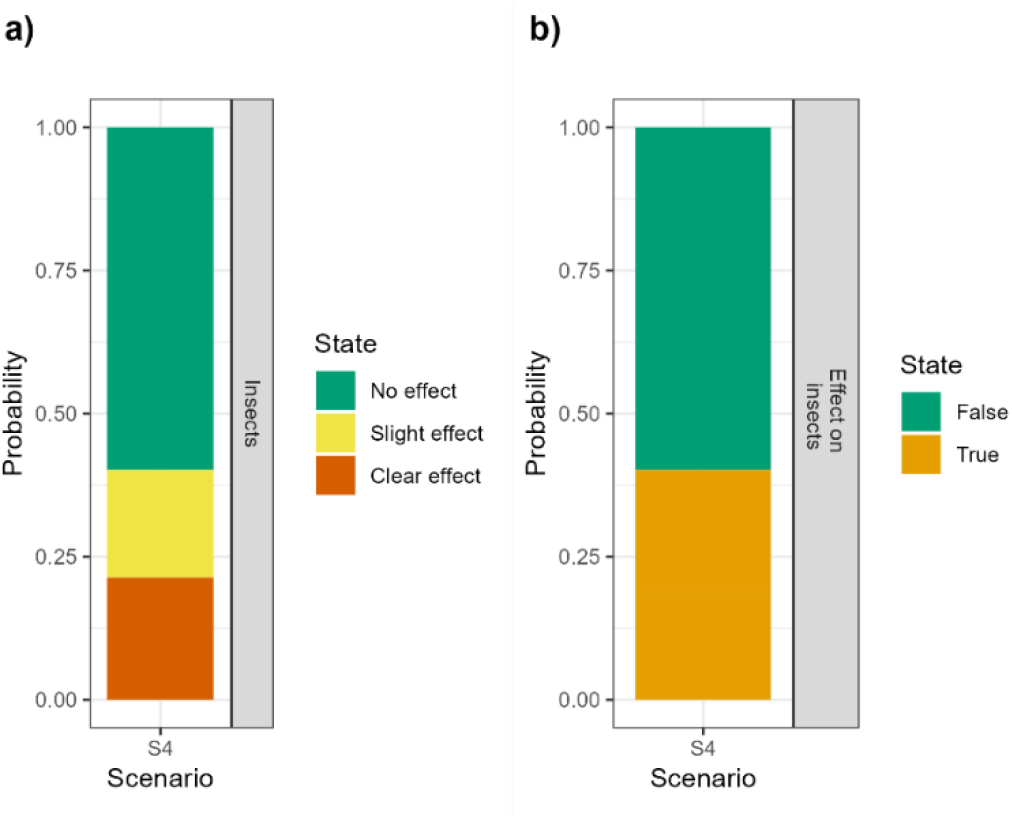
Example BN output predicted effect on insects by acetamiprid for a specific scenario (a) and summarizing Boolean node output displaying whether or not an effect of the pesticide can be assumed (b), for climate condition in 2050 and a baseline+50% application.

To summarize this example, the likelihood of there being an effect on insects was true with approx. 30% and false with approx. 70% (Figure 4b).

An assumption can be made for the effect on biological endpoints and the endpoint groups (Figure 5). When comparing some of the biological endpoints for the insecticide acetamiprid, it can be observed that macroinvertebrate were predicted to not be affected by acetamiprid with a likelihood of almost 100 %. Unlike the insects, macro-and microcrustaceans that had a higher probability to be affected, with a likelihood of 25-30%. The effect on the endpoint group coul also aggregated with the BN. In this insecticide example the biological endpoints displayed were all considered for the endpoint group of invertebrates. It can be concluded that the predicted probability of there to be an affect on any of these biological endpoints of the invertebrates endpoint group was false with approx. 25 % and true with approx. 75%. In other words, this means that it is more probable for at least one of the the bilological endpoints to be affected.

**Figure 5.**
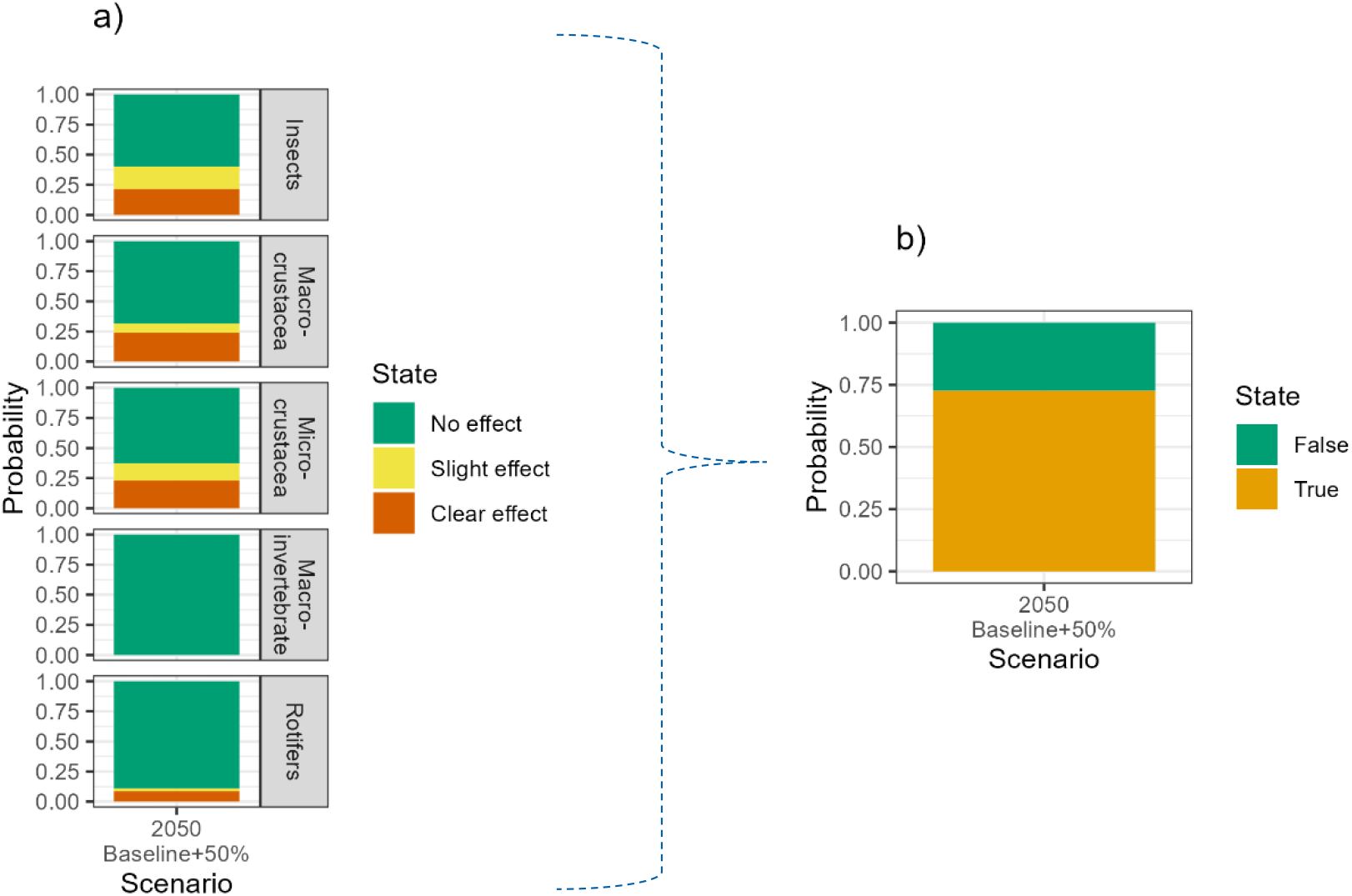
Example for the predicted effect of insecticide on the biological endpoints (a), which are considered in the endpoint functional group “Invertebrates” (b), for climate conditions in 2050 and baseline+50% application.

### 3.3. Comparison of the predicted effect of pesticides on the endpoint groups and community

Other output of the BN was the effect on the endpoint groups and community, which describes the combined probability of any of the biological endpoints being affected.

The fungicide azoxystrobin, has a probability of 50% for any of its biological endpoints being affected of the plant endpoint group (for the baseline application scenario). The endpoint group of ecosystem system processes had a probability of less than 50% for any of its biological endpoints to be affected. On the other hand, the vertebrates were not affected (Supplement Information I Figure S. 8 – upper panel).

For the herbicide MCPA, the probability of any of the plant’s biological endpoints being affected was about 50%. With a probability of 15-20% of any of the biological endpoints of the ecosystem processes were affected by the herbicide. Again, no biological endpoint was affected for the vertebrate endpoint group. (Supplement Information I Figure S. 10 – lower panel).

With a probability of 50-75% it was predicted that any of the biological endpoints of the invertebrates were affected by the insecticide acetamiprid. In contrast, the endpoint groups of plants and ecosystem process were predicted to have mostly none of the biological endpoints affected. For this pesticide type, the vertebrates were expected to not be affected (Supplement Information I Figure S. 6 – lower panel).

The summarizing effect on community node shows that all three pesticides influenced at least one of the biological endpoints (Figure 6). The fungicide (Azoxystrobin) affected any of the biological endpoints of the community with a predicted probability of almost 100%, while the herbicide (MCPA) had almost 90 % probability of affecting one of the endpoints. The insecticide (Acetamiprid) had the lowest predicted risk to the community, with a probability of almost 75% of any of the biological endpoints being affected.

**Figure 6.**
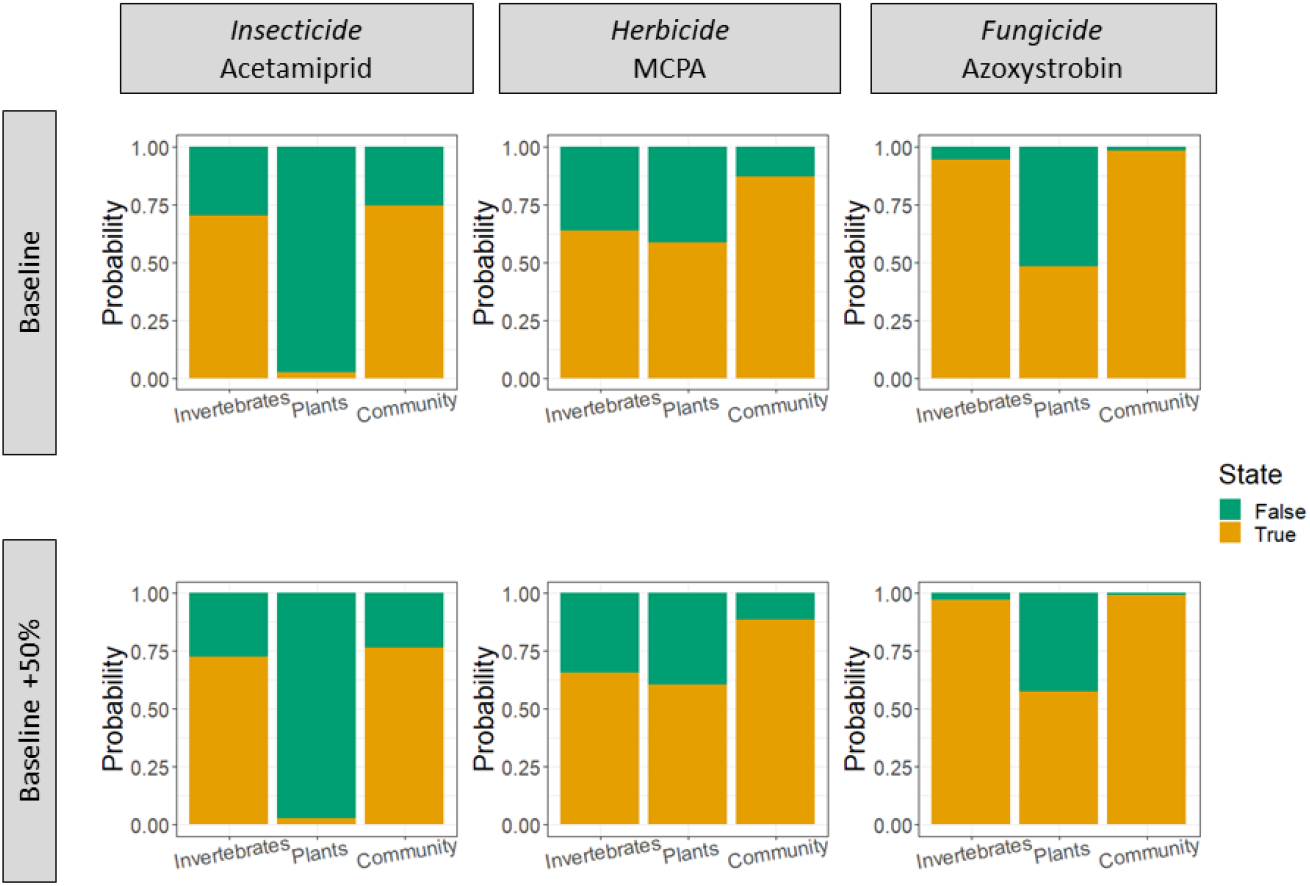
Example of effect on invertebrates, plant endpoint group and community displaying the probability in a bar chart for the selected pesticides, for a baseline and baseline+50% application under climate conditions in 2050.

In the following example, the results display the effect on any of the biological endpoints of the invertebrate, plant endpoint groups and community for either baseline and baseline+50% application scenario under the same future climate conditions (2050) (Figure 6). Focusing on the endpoint groups, the invertebrates had highest probability of being affected by azoxystrobin and lowest by acetamiprid. It was observed that the probability of an effect on any of the biological endpoints of the plant community (endpoint group) was highest for MCPA with approx. 60 % (for the baseline application), and lowest for acetamiprid with approx. 2%. Azoxystrobin has the highest probability (98%) of any of the biological endpoints in the community being affected, and Acetamiprid had the lowest probability with 75%. From these BN predictions, it can also be observed that the increase in effect of any of the biological endpoint is highest for azoxystrobin whenever a higher application scenario is used, whereas the lowest increase in probability was observed for acetamiprid.

## 4 Discussion

In general, under any of the scenario combinations the BN predicted at least one of the biological endpoints were effected by the pesticicdes. Azoxystrobin was most likely to effect at least one of the biological endpoints, followed by MCPA that also had a high likelihood, whereas Acetamiprid compared to the other two pesticides had the lowest. Furthermore, a trend from the predictions could be observed for the higher application of pesticides that resulted in a shift towards a higher probability for any of the biological endpoint and endpoint groups being affected for all pesticides. On the other hand, the climate condition at the time had no trend on the effect on biological endpoints that can be observed for all pesticide types.

We aimed to carry out effect assessment of three selected pesticides using a probabilistic approach, as these have been recommended to better account for uncertainty in pesticide exposure (Carriger & Newman, 2012) and effect (Dreier et al., 2020). Therefore, we linked the inputs and outputs of two prediction models into a Bayesian network (BN). We had succeded in developing a BN model that can predict the effect on multiple biological endpoints, as well as cumulative effect on endpoint groups and community. These BN predictions enabled comparison between different pesticide types, community levels, as well as pesticide application and climate change scenarios.

Some initial precision of the prediction model outputs might be lost due to a common BN shortcoming when discretization of continuous variables is applied (Marcot, 2017; Nojavan et al., 2017). A higher resolution of BN predictions can be achieved by applying dynamic discretization (Carriger et al., 2016; Fenton & Neil, 2018). The credibility of developed Bayesian networks is mainly influenced by the assumptions and input data derived from predictions derived with the process-base exposure model and case-based effect model. The RICEWQ model used to predict the exposure concentrations in the rice paddy is readily available for simulation and can be used for higher tier exposure assessment (MED-Rice, 2003). A detailed description of uncertainties related to this model can be found in Miao et al. (2004). We chose this model as it enabled simulation of agricultural conditions for rice production such as controlled release of water, overflow, and flooding unlike other pesticide fate and transport models. Moreover, it was considered as the first option when carrying out an exposure prediction for rice cultures (MED-Rice, 2003). However, there were some assumptions and input data that lead to uncertainty in our modelling efforts. We could have derived more realistic model outputs by updating or adding scenarios to our model efforts. The use of multiple climate models with different greenhouse gas emission scenarios allows integration of more variability (Fernández et al., 2017). The model could he been improvement by using more and different climate models as some papers by Steffens et al. (2014) and Moe et al. (2022) recommended. These mentioned that using an ensemble of various lobal and regional climate models together with various greenhouse gas emission scenarios would potentially enable more robust estimations of pesticide losses in future. In addition, we assumed more realistic application scenarios could be developed and used to run the autoRICWQ model. Future work for the prediction of the exposure concentration could also be extended to the discharge channel, by using the RIVWQ model, which would enable the consideration and integration of dilution better.

There is also some uncertainty associated with the PERPEST model, connected to the ecotoxicology database. The database that the case-based effect model bases its predictions on has limited data availability for fish and tadpoles. Furthermore, its incorporated data is primarily based on datasets from temperate climate such as Europe and North-America (Davis et al., 2013; Van den Brink et al., 2002), and therefore is limited in its predictions for the Mediterranean climate zone. Henceforth, this limitation could be overcome by updating the database with more and regional relevant bioassays. Some other uncertainties of the PERPEST model can be associated with the input information such as pesticide properties for the model run. Consequently, we tried to minimize this, by using the same information source for the selected pesticides whenever possible. For example, we collected toxicity data from ECOTOXicology Knowledgebase (Olker et al., 2022) and thereafter using the same method to prepare the data used on MOSAIC (Charles et al., 2017) to predict the HC50. In essence, the effect prediction model has simple data requirements making it easy to use (Davis et al., 2013). Thereupon, some uncertainty is also linked to the way the PERPEST model output is integrated in the BN, as it is a gradient and its concentration range thus far cannot be adjusted to fit better with the exposure distribution. Some other restrains are pointed out by Davis et al. (2013) detailing that PERPEST output might be challenging to use and understand by stakeholders and used in as risk management due to the lack of a “established threshold risk value”. To overcome this limitation of the PERPEST model Davis et al. (2013) suggested to set acceptable probabilities. In this study, we tried a different approach to enable easy communication of BN outputs integrated a summarizing node for the effect on endpoint group and community. These nodes show the probability of any of the biological endpoints to be affected to be true or false. This far there is no direct link from the scenarios to the effect module of the network. This relationship needs to be further explored, as the combined effect of climate conditions and chemical exposure are expected to change the effect on the different biological endpoints.

Additional research and model development may result in a better integration and use of the prediction model outputs. An updated PERPEST model database would greatly decrease uncertainties. When it comes to the RICEWQ model calibration larger number of models runs with more application and climate scenarios, and crop types would be beneficial to increase reflection of variability. In addition, the BN model could also be run for other pesticides commonly used for rice production in the area. Thenceforward this could also allow the prediction of cumulative risks of intentional mixtures. Most of these improvements are likely to require some changes to the model structure though. Nevertheless, this model enables accounting for uncertainty of all compartments of the BN model, which allows for transparency when communication the effect of pesticides to various biological endpoints, endpoint groups and the community in the aquatic environment.

## 5 Conclusion and future outlook

This study shows how to use a Bayesian network model to integrate the outcomes of two different predictive models - the pesticide exposure model RICEWQ and the biological effect model PERPEST, and thereby predicts the risk of a pesticide on biological endpoints and endpoint groups in the aquatic ecosystem of a rice paddy. The BN, we have developed can carry out probabilistic calculation of risk for various event such as pesticide application scenarios. This approach builds upon our probabilistic model versions of the traditionally used Risk Quotient calculation, displaying uncertainty transparently of all its model components. The current study further expands this approach by including the risk calculation for individual biological endpoints as well cumulative risk for the endpoint groups and the community.

For example, the fungicide azoxystrobin was predicted to have the highest probability (about 98%) of affecting any of the biological endpoints in the community. Followed by the herbicide MCPA, which had a probability of 85% of affecting any of the biological endpoints in the community. MCPA, compared to azoxystrobin, the invertebrate’s endpoint group had lower probability of any of the endpoints being affected, and the plant endpoint group a higher probability. The insecticide Acetamiprid had the lowest probability of affecting any of the biological endpoint groups in the community. In comparison to the two other pesticides, its plant community (endpoint group) has higher probability of none of the biological endpoints to be affected.

Future research efforts can to incorporate more scenarios such as additional crop types, application patterns and an ensemble of climate models to derive a more realistic idea of pesticide effects on the aquatic ecosystem in future. In addition, we aim to carry out an effect assessment of the intentional mixtures applied in the Albufera national park, to move away from single compound assessment.

## Supporting information

Supplement Information III

## Acknowledgement

This research was funded by ECORISK2050, which has received funding from European Union’s Horizon 2020 research and innovation program under the grant agreement No. 813124 (H2020-MSCA-ITN-2018). As well as, the CICLIC project, funded by the Spanish Ministry of Science, Innovation and Universities (RTI 2018_097158_A_C32), and the Talented Researcher Support Programme - Plan GenT (CIDEGENT/2020/043) of the Generalitat Valenciana. K. E. Tollefsen was funded by NIVA’s Computational Toxicology Program (www.niva.no/nctp).

We thank Yasser Fuentes-Edfuf for his help in probabilistic exposure modelling. We thank Jes Rasmussen (NIVA), discussion and advice on the pesticide effect modelling. As well as Wayne Landis (Western Washington University) and John Carriger (USEPA) for advice on Bayesian network modelling.

## Software availability

RICEQW - Please contact to register to receive a download of this model. autoRICEWQ code - https://zenodo.org/record/5940235#.YhXv8uiZMZA

A free version of NeticaTM is available online at: http://www.norsys.com/downloads.html, along with a glossary of Bayesian network terms and tutorials.

## List of figures in Supplement

Figure S. 1 Example of the predicted effect on the taxonomic group Algae and Macrophytes that where derived for the herbicide.

Figure S. 2 Bayesian network for the insecticide

Figure S. 3 Bayesian network for the fungicide

Figure S. 4 Bayesian network for the herbicide

Figure S. 5 Overview of effects on the biological endpoints by the selected insecticide (acetamiprid) for all selected scenarios.

Figure S. 6 Overview of effects on the endpoint groups community level by the selected insecticide (acetamiprid) for all selected scenarios.

Figure S. 7 Overview of effects on the biological endpoint by the selected fungicide (azoxystrobin) for all selected scenarios.

Figure S. 8 Overview of effects on the endpoint groups community level by the selected fungicide (azoxystrobin) for all selected scenarios.

Figure S. 9 Overview of effects on the biological endpoint by the selected herbicide (MCPA) for all selected scenarios.

Figure S. 10 Overview of effects on the endpoint groups and community level by the selected herbicide (MCPA) for all selected scenarios.

## Supplement information overview

- Overview PERPEST model inputs
- Overview autoRICEWQ inputs
- Overview of BN output

