## Supplement Information I for "Using a Bayesian network model to predict effects of pesticides on aquatic community endpoints in a rice field – A southern European case study"

### Supplement Information I – RICEQW & PERPEST Paper

#### 1. Endpoint groups per pesticide type

The following lists the biological endpoint groups that are considered by the PERPEST model.

**Fungicide:**

- Invertebrates: Microcrustacea, Macrocrustacea, Insecta, Other zooplankton taxa, Other macro-invertebrate taxa
- Plant: Periphytic algae, Phytoplankton, Macrophytes
- Vertebrates: Fish and tadpoles
- ecosystem process: DO-pH metabolism, Decomposition

**Insecticide:**

- vertebrates: Fish
- invertebrates: Insects, Macrocrustacea, Microcrustacea, Other macro-invertebrates, Rotifers
- plants: Algae and macrophytes
- ecosystem process: Community metabolism

**Herbicide:**

- ecosystem process: Community metabolism
- invertebrates: Zooplankton, Macrocrustaceans & Insects, Molluscs
- vertebrates: Fish and Tadpoles
- plants: Macrophytes, Periphyton, Phytoplankton

#### 2.  PERPEST model description and output

Description of assumption and processes on the PERPEST model

In contrast to most effect models, PERPEST is based on empirical data from micro- and mesocosms extracted from literature. It searches for situations in the database which resemble the case in question, based on relevant (toxicity) characteristics of the compound. This allows the model to predict effects of pesticides for which no evaluation on a semi-field scale have been published. PERPEST results in a prediction showing the probability of three effect classes (no, slight or clear effects) for the various grouped endpoints. For each taxon group, the predicted probability of effect classes along the pesticide concentration gradient is used to derive the conditional probability table for this taxon node in the BN.

Example gradient output for an insecticide Figure S. 1.

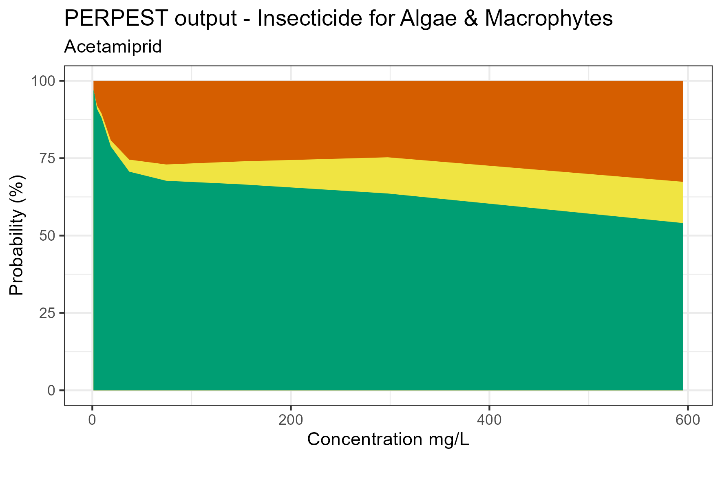

Figure S. 1 Example of the predicted effect on the taxonomic group Algae and Macrophytes that where derived for the herbicide.

#### 3. Bayesian network output for the biological endpoints and endpoint groups

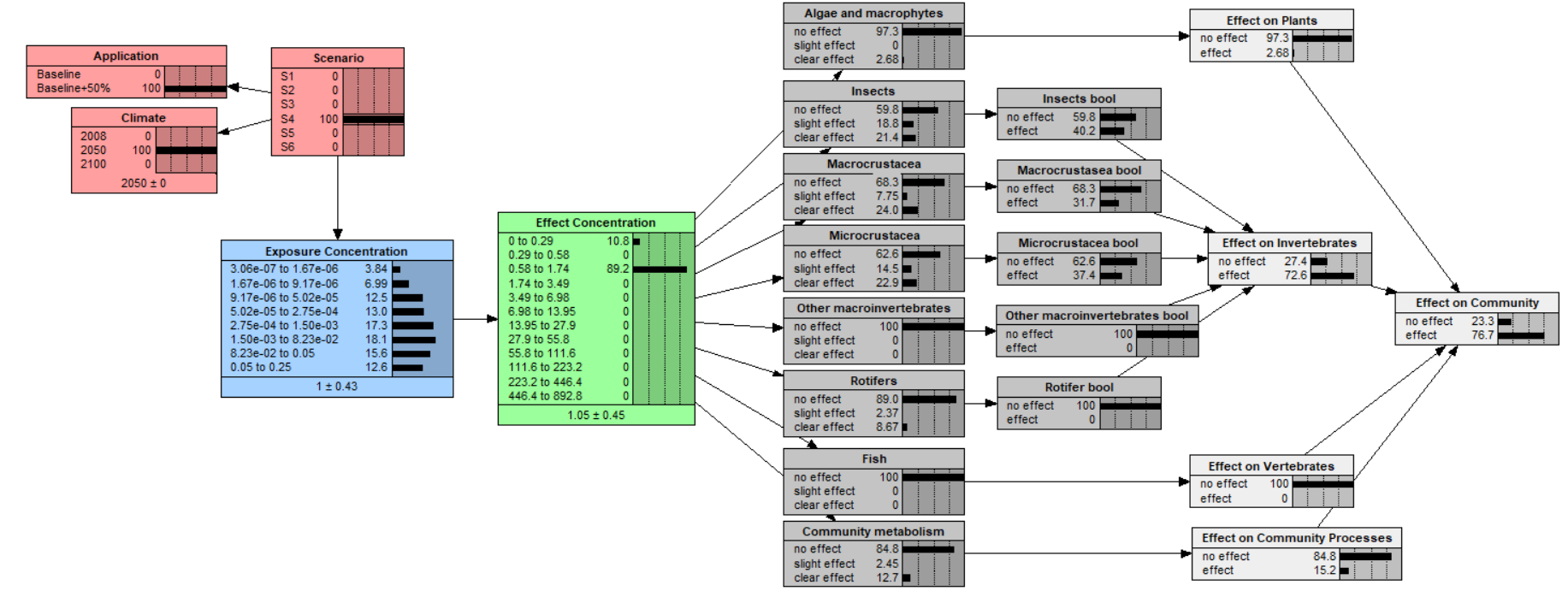

Figure S. 2 Bayesian network model for the insecticide

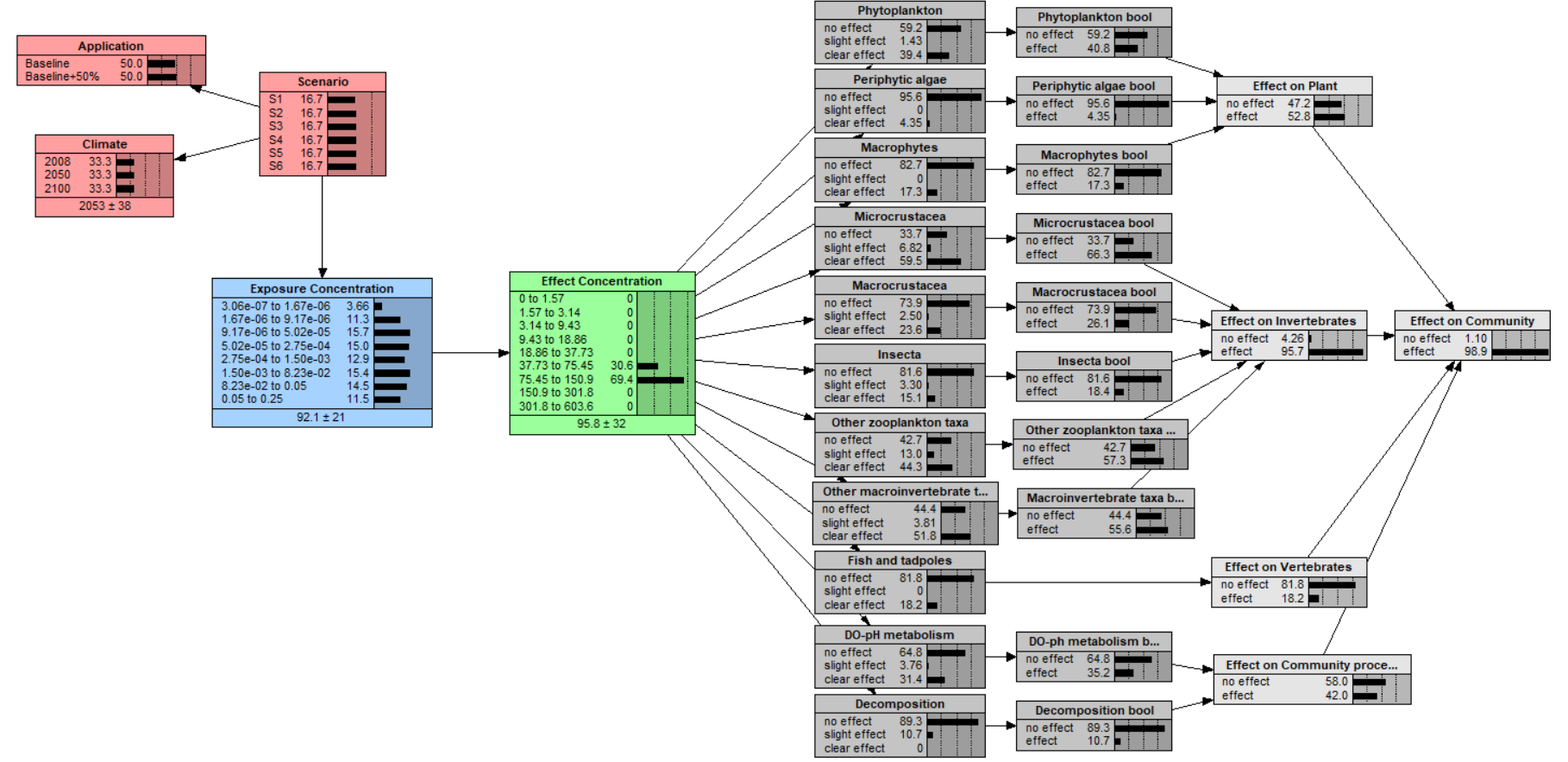

Figure S. 3 Bayesian network for the fungicide

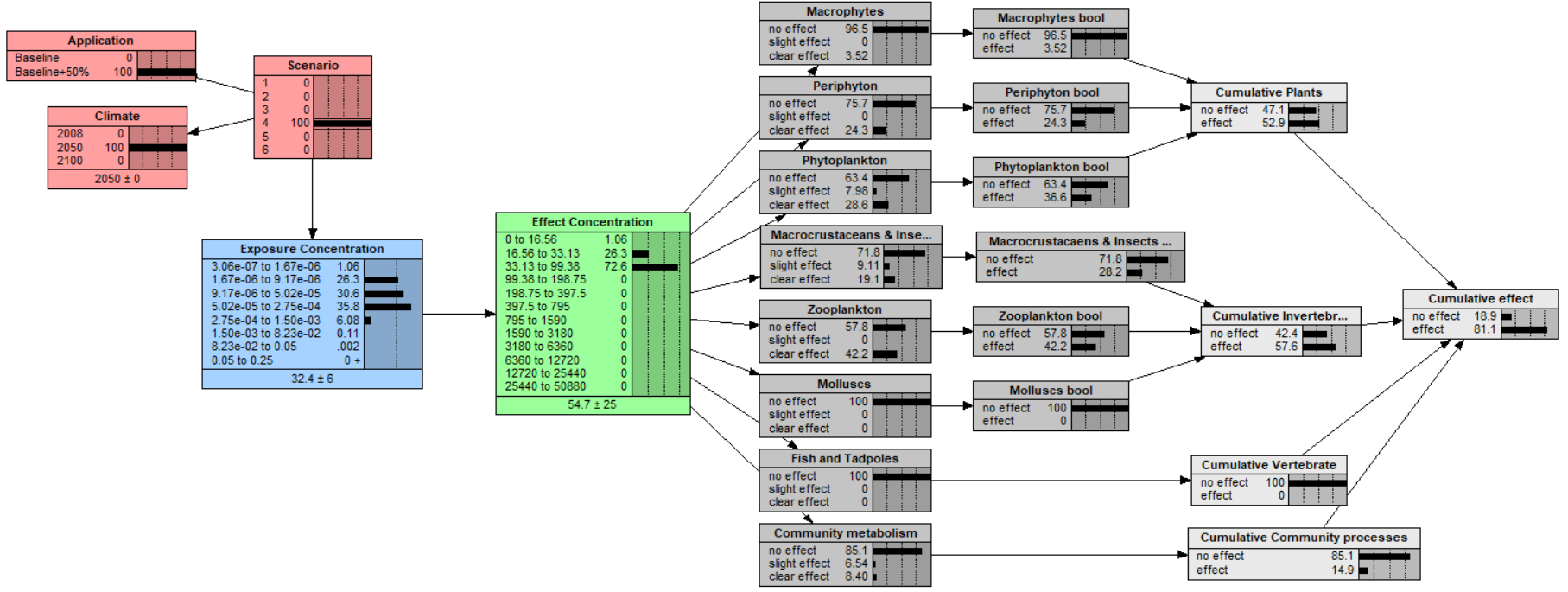

Figure S. 4 Bayesian network for the herbicide

It can be observed that for the insecticide. The probability of the biological endpoints to had slight and clear effect state was higher in 2050 and 2100. The opposite could be observed for the fungicide azoxystrobin, here the probability of the biological endpoint to be in the slight and clear effect state decrease in 2050 and 2100. The herbicide MCPA, also showed a decrease of probability of the biological endpoints to be in slight and clear effect state for 2100 and predicted a similar probability of the previous climate-time scenarios. For the fungicide, the probabilities for any of the biological endpoints in the endpoint groups didn’t seem to change much over the years, with one exception, ecosystem processes had a higher probability of none of the biological endpoints to be affect in 2100. As for the herbicide, in general azoxystrobin had lower effect on the endpoint groups in 2100 than previous time-periods. The overall trend for the insecticide was that the likelihood of any of the biological endpoints not being affected was highest in 2008 and lowest in 2050. The Probability were slightly higher for the biological endpoints to be affected by MCPA in 2050 and 2100. Whereas, a decreased was observed for azoxystrobin for the climate conditions in 2050 and 2100. For acetamiprid an increase was observed in probability for a biological endpoint to be affected for the climate conditions in 2050 for acetamiprid followed by a decrease in 2100.

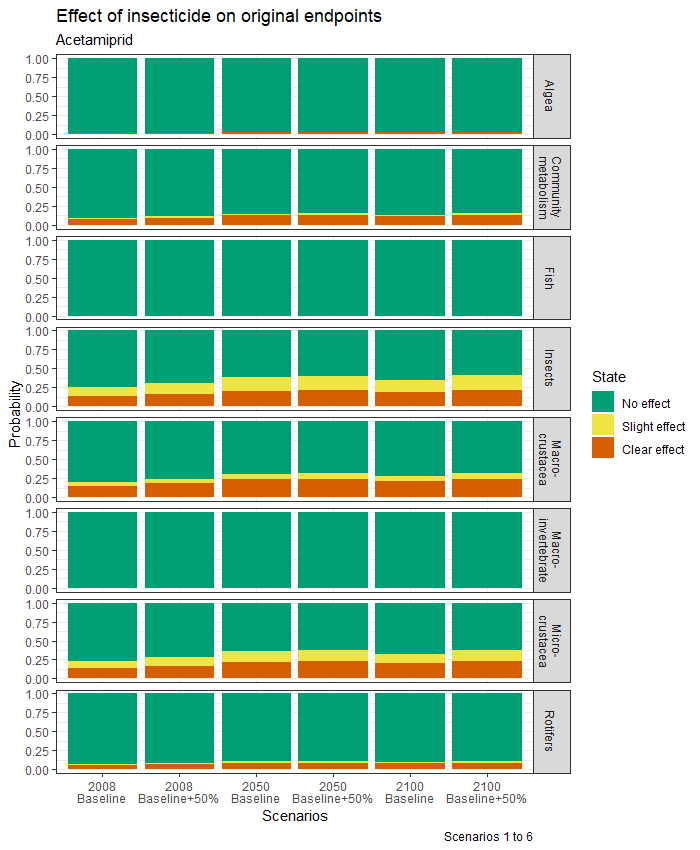

Figure S. 5 Overview of effects on the biological endpoints by the selected insecticide (acetamiprid) for all selected scenarios.

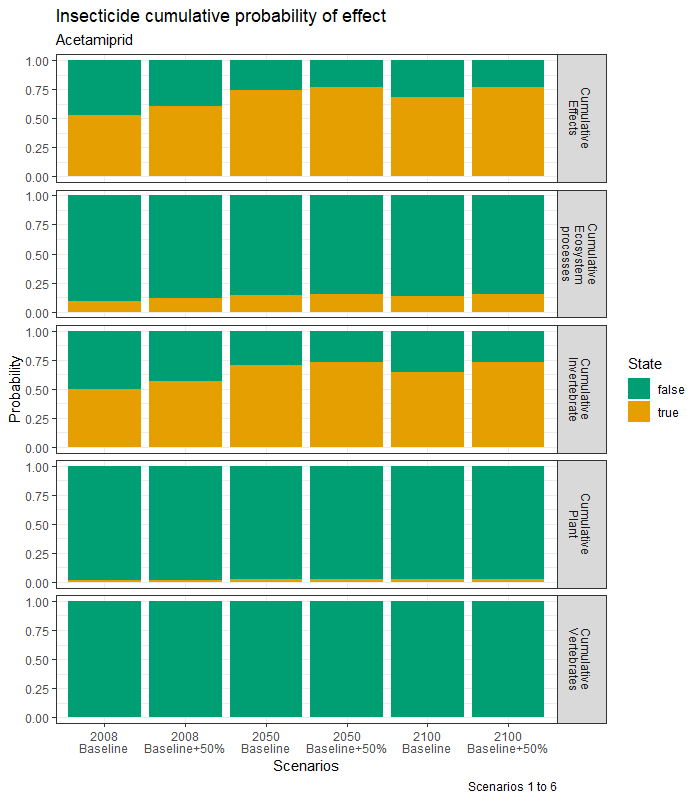

Figure S. 6 Overview of effects on the endpoint groups community level by the selected insecticide (acetamiprid) for all selected scenarios.

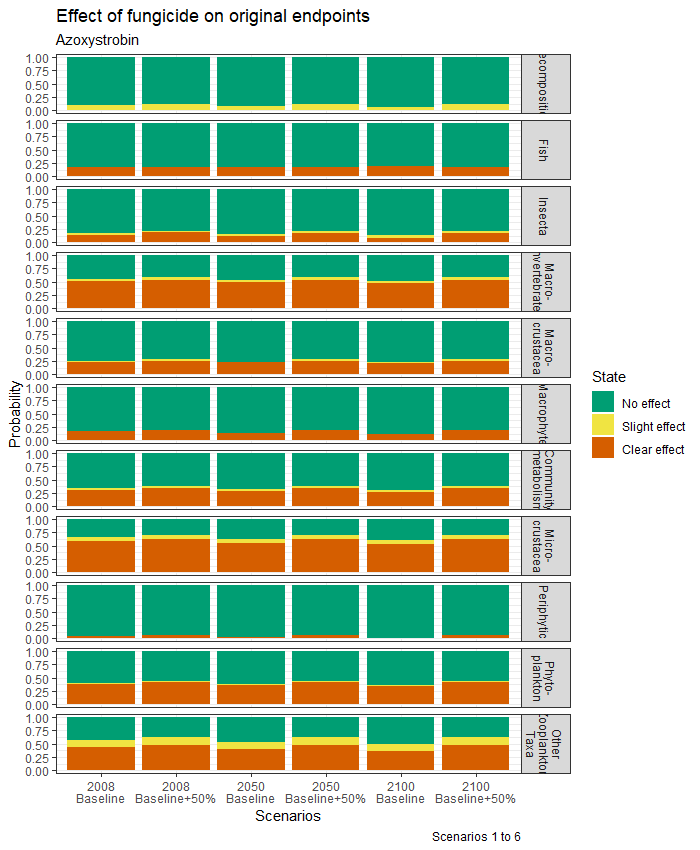

Figure S. 7 Overview of effects on the biological endpoint by the selected fungicide (azoxystrobin) for all selected scenarios.

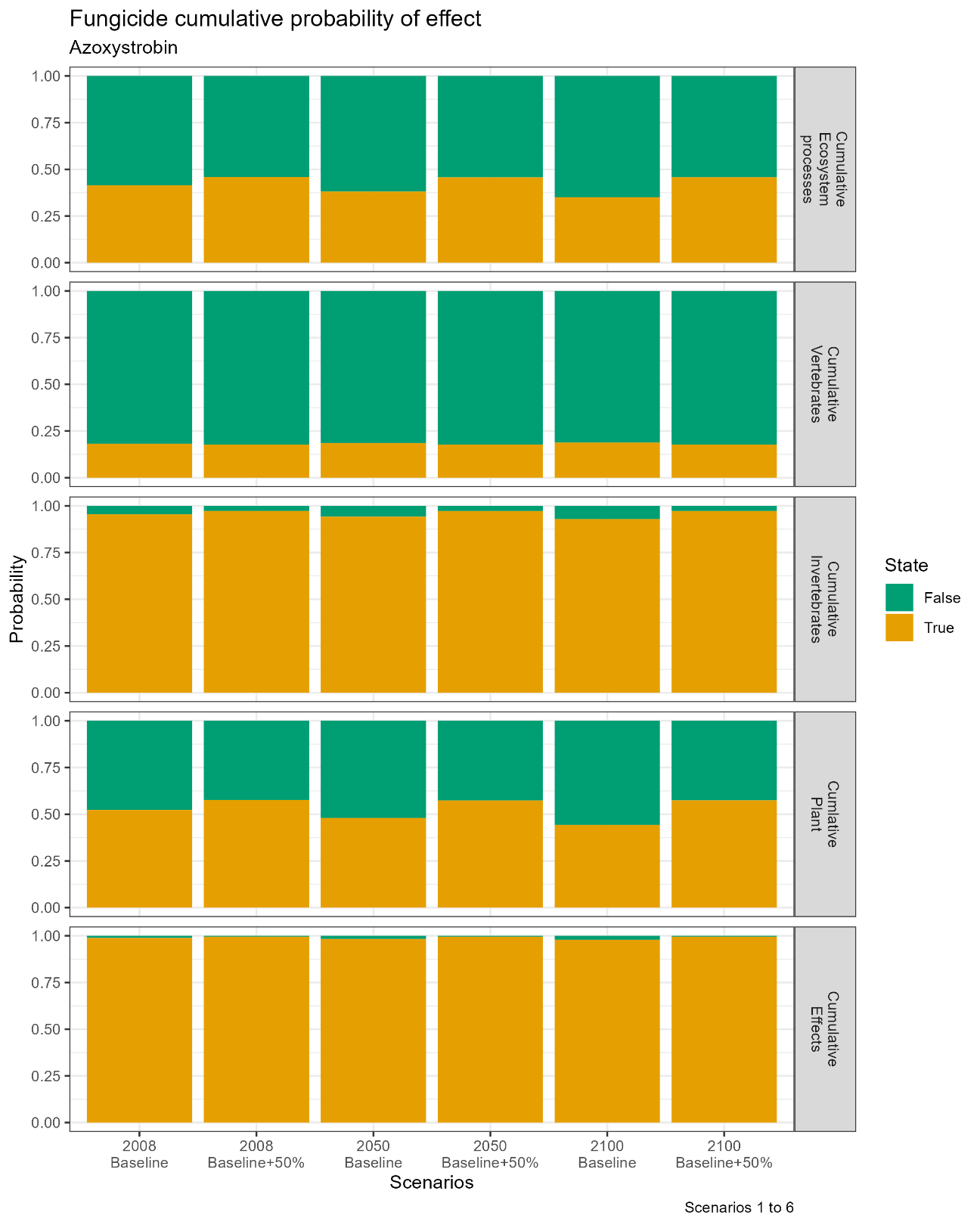

Figure S. 8 Overview of effects on the endpoint groups community level by the selected fungicide (azoxystrobin) for all selected scenarios.

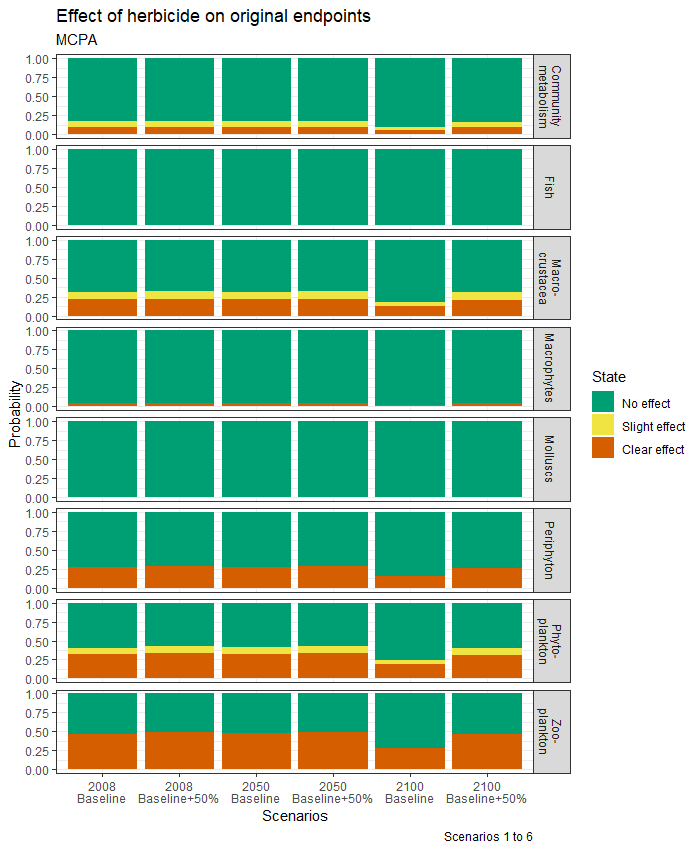

Figure S. 9 Overview of effects on the biological endpoint by the selected herbicide (MCPA) for all selected scenarios.

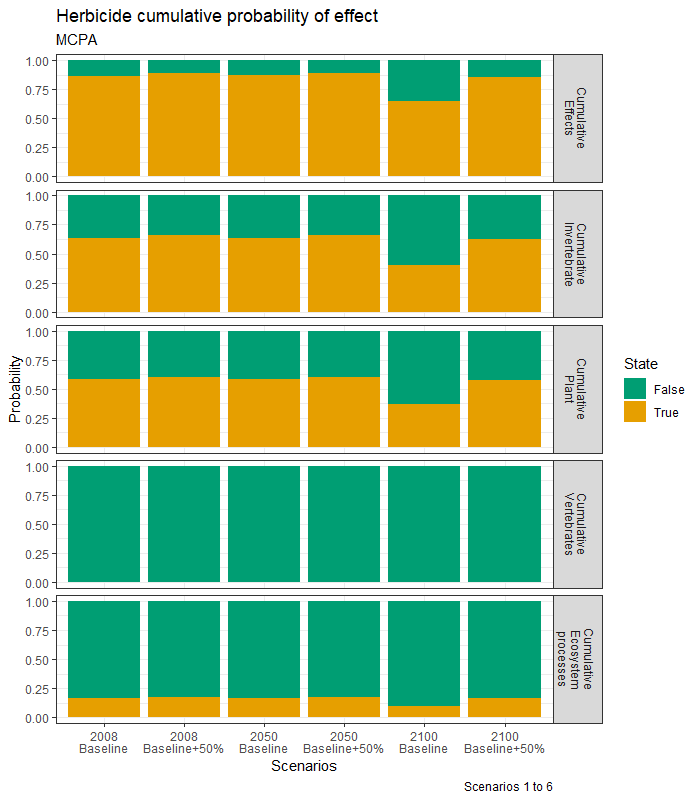

Figure S. 10 Overview of effects on the endpoint groups and community level by the selected herbicide (MCPA) for all selected scenarios.
